# A single amino acid in the *Salmonella* effector SarA/SteE triggers supraphysiological activation of STAT3 for anti-inflammatory target gene expression

**DOI:** 10.1101/2024.02.14.580367

**Authors:** Margaret R. Gaggioli, Angela G. Jones, Ioanna Panagi, Erica J. Washington, Rachel E. Loney, Janina H. Muench, Richard G. Brennan, Teresa L. M. Thurston, Dennis C. Ko

**Affiliations:** Department of Molecular Genetics and Microbiology, School of Medicine, Duke University, Durham, NC 27710, USA; Department of Infectious Disease, Centre for Bacterial Resistance Biology, Imperial College London, London, UK; Department of Biochemistry, School of Medicine, Duke University, Durham, NC 27710, USA; The Francis Crick Institute, London NW1 1AT, UK; Division of Infectious Diseases, Department of Medicine, School of Medicine, Duke University, Durham, NC 27710, USA

**Keywords:** STM2585, gogC, JAK-STAT, SOCS3, IL6ST, Y705, IL-6

## Abstract

Non-typhoidal *Salmonella enterica* cause an estimated 1 million cases of gastroenteritis annually in the United States. These serovars use secreted protein effectors to mimic and reprogram host cellular functions. We previously discovered that the secreted effector SarA (*Salmonella* anti-inflammatory response activator; also known as SteE) was required for increased intracellular replication of *S.* Typhimurium and production of the anti-inflammatory cytokine interleukin-10 (IL-10). SarA facilitates phosphorylation of STAT3 through a region of homology with the host cytokine receptor gp130. Here, we demonstrate that a single amino acid difference between SarA and gp130 is critical for the anti-inflammatory bias of SarA-STAT3 signaling. An isoleucine at the pY+1 position of the YxxQ motif in SarA (which binds the SH2 domain in STAT3) causes increased STAT3 phosphorylation and expression of anti-inflammatory target genes. This isoleucine, completely conserved in ∼4000 *Salmonella* isolates, renders SarA a better substrate for tyrosine phosphorylation by GSK-3. GSK-3 is canonically a serine/threonine kinase that nonetheless undergoes tyrosine autophosphorylation at a motif that has an invariant isoleucine at the pY+1 position. Our results provide a molecular basis for how a *Salmonella* secreted effector achieves supraphysiological levels of STAT3 activation to control host genes during infection.

## Introduction

*Salmonella enterica* serovar Typhimurium is a gram-negative facultative intracellular anaerobe and a causative agent of salmonellosis, a gastrointestinal infection that is one of the most common foodborne diseases in humans. Salmonellosis outbreaks are a major public health concern, with an estimated 1 million annual cases in the United States alone.^1^

Essential to the pathogenesis of *S.* Typhimurium – and all *Salmonella enterica* serovars – is an arsenal of secreted protein effectors.^2^ These effectors are translocated into the host cell by type III secretion systems (T3SS) encoded within *<u>S</u>almonella* <u>p</u>athogenicity islands (SPI).^3^ Secreted protein effectors can mimic and reprogram host cellular functions to create a beneficial environment for the invading bacteria, such as formation of the *Salmonella-*containing vacuole (SCV) and antagonization of the immune response.^4^ While secreted factors are important to all *Salmonella* serovars, different serovars have different repertoires of effectors, reflecting the diverse niches and pathogenic outcomes seen with *S. enterica*.

We previously used this natural diversity to identify the effector SarA (Salmonella anti-inflammatory response activator; also known as Stm2585, GogC, and SteE). SarA acts through activation of the transcription factor STAT3 to regulate the host response to *S.* Typhimurium infection.^5^ SarA is secreted primarily by the SPI-2 T3SS ^5^ and recruits the kinase GSK-3 to phosphorylate STAT3 at Y705,^6,7^ stimulating intracellular bacterial replication and production of the anti-inflammatory cytokine interleukin-10 (IL-10),^5^ as well as M2 polarization of macrophages.^7–9^

SarA recruits STAT3 to the GSK-3-SarA complex through an amino acid motif that is similar to the host cytokine co-receptor gp130.^6^ Paradoxically, gp130 is the shared signal transducing subunit for the pro-inflammatory IL-6 family of cytokines, yet SarA activation of STAT3 leads to production of IL-10 (which is itself a stimulator of STAT3 activation). Studies comparing pro- and anti-inflammatory STAT3 signaling pathways support the idea that regulation of the strength and duration of STAT3 activation impacts downstream transcriptional targets.^10–14^ IL-6 stimulation of human cell cultures leads to a sharp, transient increase in STAT3 phosphorylation, while more robust and prolonged phosphorylation of STAT3 has been observed with IL-10 stimulation. Furthermore, prolonging phosphorylation through deletion of negative regulators^11,12^ or use of constitutively active gp130 dimers and STAT3^13,15^ transforms the transcriptional response induced upon IL-6 stimulation to become more similar to that induced by IL-10. These results suggest that the nature of STAT3-induced transcriptional changes is dictated by the strength and duration of phosphorylation. Therefore, we hypothesized that there are molecular characteristics of SarA-directed phosphorylation of STAT3 that make it more similar to anti-inflammatory activation than pro-inflammatory activation.

Here, we compared STAT3 activation mediated by SarA vs. gp130 to understand the nature and molecular basis of SarA-induced activation on the host transcriptional response. In response to SarA transfection or during infection, we observed greater phosphorylation of STAT3 and upregulation of anti-inflammatory genes. We hypothesized that more robust and prolonged activation by SarA could be due to either increased STAT3 binding affinity or a lack of the negative regulation that canonically dampens gp130 signaling. Our results demonstrate that the amino acid residue at the pY+1 position of the SarA STAT3-binding YxxQ motif is critical for the increased binding of STAT3 to SarA compared to gp130. Unexpectedly, this effect is mediated through increased tyrosine-phosphorylation of the SarA YxxQ motif by GSK-3, rather than an increased binding affinity to STAT3. The isoleucine at this position in SarA is invariant across 4,554 *S. enterica* sequences and matches the isoleucine conserved across 245 homologs in GSK-3’s autophosphorylated site (which has the sequence YICS). Thus, while SarA shares a 39-amino acid region of sequence similarity with gp130, SarA promotes supraphysiological activation of STAT3 because of a single amino acid difference that matches the substrate specificity of GSK-3, a host kinase with high baseline activity.

## Results

### SarA activation of STAT3 demonstrates an anti-inflammatory bias

The STAT3 signaling pathway can be activated by a wide variety of signals leading to either pro- or anti-inflammatory transcriptional responses. SarA was named because of its induction of IL-10 and other anti-inflammatory genes, but we also observed induction of pro-inflammatory genes such as *IFNG* and *CXCL9* in our previously published transcriptomics dataset comparing LCLs (lymphoblastoid cell lines, immortalized B cells) infected with wild type or *ΔsarA S.* Typhimurium.^5^ Therefore, to better characterize the nature of SarA-dependent activation of STAT3, we compared our transcriptomics data to publicly available data generated from human monocyte-derived dendritic cells stimulated with either IL-6 or IL-10.^10^ We determined that genes activated by SarA during S. Typhimurium infection are significantly enriched for IL-10 activated genes (p=2.2e-16; 3.53-fold enrichment). In contrast, no enrichment is noted for IL-6 activated genes (p=1) (**Fig. 1A**). Canonical anti-inflammatory targets like *TNIP3,* which is a negative regulators of NF-κB signaling, are upregulated in both a SarA and IL-10 dependent manner. *SOCS3* and *SBNO2*, which are known negative regulators of the IL-6 signaling pathway, are significantly upregulated in all three conditions, but are induced to a greater extent during IL-10 stimulation as compared to IL-6 (**Fig. S1**). Thus, there are both qualitative and quantitative differences in STAT3 targets, depending on the stimulus, and the greater overlap of SarA and IL-10 transcriptional targets demonstrates that SarA-dependent activation of STAT3 has an anti-inflammatory bias.

**Figure 1:**
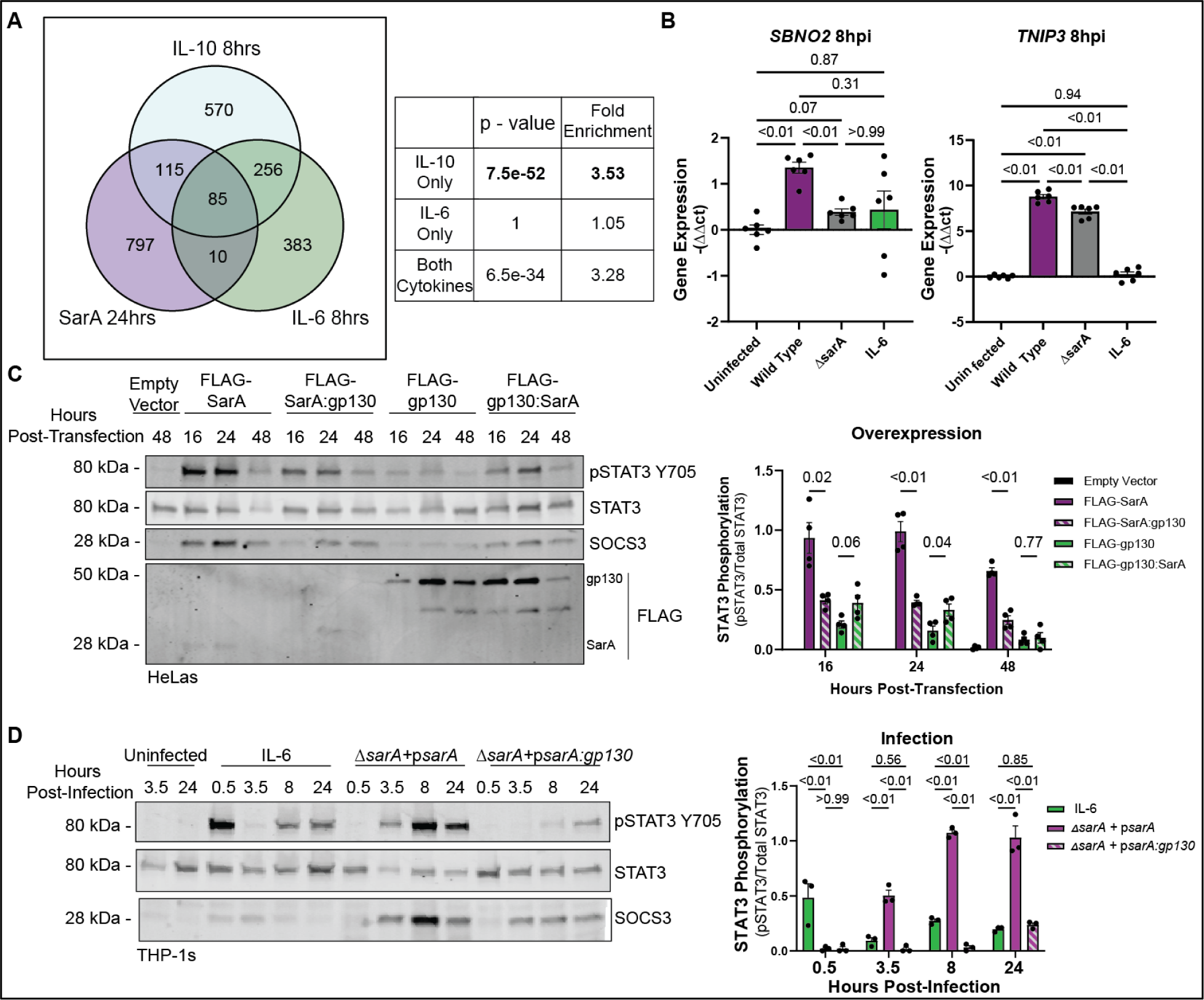
SarA induces anti-inflammatory cell signaling and strong STAT3 phosphorylation. (A) SarA upregulated genes significantly overlap with IL-10 target genes. Venn diagrams showing overlap of genes upregulated in a SarA-dependent manner (during *S.* Typhimurium infection in LCLs from ^5^) and in an IL-10 or IL-6 dependent manner (after cytokine stimulation in dendritic cells from ^10^). P-values obtained from a chi-squared test. (B) Wild-type *S.* Typhimurium infection leads to increased expression of anti-inflammatory genes. THP-1 cells were either stimulated with 10 ng/mL of IL-6 or infected with wild-type or Δ*sarA S.* Typhimurium. RNA was collected from cells 8 hrs post-infection and expression levels of *SBNO2* and *TNIP3* were measured via qPCR. Data are from three experiments. P-values obtained from a Brown-Forsythe and Welch ANOVA with Dunnett’s T3 multiple comparisons test. (C) SarA GBS domain leads to greater STAT3 phosphorylation compared to gp130 GBS domain as quantified by western blot after overexpression in HeLa cells. Data are from four experiments. P-values obtained from unpaired t-tests with Welch’s correction comparing FLAG-SarA to FLAG-SarA:gp130 and FLAG-gp130 to FLAG-gp130:SarA. (D) SarA GBS domain leads to greater STAT3 phosphorylation compared to gp130 GBS domain as quantified by western blot after *S.* Typhimurium infection in THP-1 cells. Data are from four experiments. P-values obtained from a 2-way ANOVA with Tukey’s multiple comparisons test.

To confirm this SarA-dependent induction of anti-inflammatory targets during infection, we measured the expression levels of anti-inflammatory targets identified from our comparative transcriptomics analysis during wild-type or *ΔsarA S.* Typhimurium infection (**Fig. 1B**). In THP-1 monocytes, wild-type infection leads to significantly higher transcript levels of *SBNO2* and *TNIP3* compared to *ΔsarA.* The induction of these anti-inflammatory genes is not simply secondary to IL-10 production, as THP-1 monocytes demonstrated minimal phospho-STAT3 in response to IL-10 stimulation (**Fig. S2**).

Our data demonstrates that SarA secretion by *S.* Typhimurium during infection promotes an anti-inflammatory response. However, the molecular determinants for this anti-inflammatory bias of SarA are unknown.

### SarA induces more robust and prolonged phosphorylation of STAT3 than gp130 activation

It has previously been demonstrated that the anti-inflammatory bias of IL-10 activation of STAT3 is due to its prolonged and robust phosphorylation of STAT3.^12^ IL-6 leads to a shorter burst of STAT3 phosphorylation compared to IL-10. However, IL-6 signaling can be altered to be anti-inflammatory by prolonging STAT3 activation through deletion of negative regulators such as SOCS3 or use of constitutively active STAT3 or gp130.^11,13^ Therefore, we performed time-course experiments to test whether SarA activation of STAT3 results in a more prolonged and robust STAT3 activation compared to IL-6/gp130 activation.

We previously showed that SarA leads to the activation of STAT3 through a 39 amino acid region of homology with the intracellular portion of the cytokine co-receptor gp130, referred to as the GBS (<u>g</u>p130 <u>b</u>inding of <u>S</u>TAT3) domain.^6^ SarA mimics gp130, using a phosphorylated YxxQ motif in the GBS to bind to the STAT3 SH2 domain.^6^ Upon recruitment to the GSK-3-SarA complex, the kinase GSK-3 phosphorylates STAT3 at tyrosine residue 705 (Y705), thereby activating STAT3.^7^ We confirmed that the GBS domains of SarA and gp130 are functionally interchangeable, as demonstrated in a time course measuring the ratio of phosphorylated-STAT3-Y705 to total STAT3 protein levels in HeLa cells after overexpressing FLAG-tagged chimeric constructs where the GBS domain of SarA and gp130 had been swapped (**Fig. 1C**). While overexpression of all constructs led to STAT3 phosphorylation, constructs containing the SarA GBS domain triggered greater phosphorylation than constructs containing the gp130 GBS at all time points tested. Probing for FLAG shows the chimeric proteins are expressed at similar levels to their wild-type counterparts, so the decrease in STAT3 phosphorylation is not caused by reduced expression (**Fig. 1C**). Thus, overexpression experiments indicate that the SarA GBS results in greater levels of STAT3 phosphorylation than the homologous region from gp130 at multiple time points.

Similarly, we observed greater STAT3 phosphorylation when comparing *S.* Typhimurium infection with IL-6 stimulation (which signals through gp130 (*IL6ST*) and IL6R heterodimer) in THP-1 monocytes. IL-6 stimulation resulted in an early peak at 30 minutes. In contrast, SarA activation of STAT3 during infection led to high levels of phosphorylation by 3.5 hrs that were maintained through 24 hrs (**Fig. 1D**). During infection with a mutant *S.* Typhimurium strain where the GBS of SarA was replaced with the GBS from gp130 (p*sarA:gp130*), the level of phosphorylation never reached the level seen with wild-type SarA and was more similar to the level seen with IL-6 stimulation (**Fig. 1D**).

Next, we wanted to identify the key molecular differences between the gp130 and SarA GBS domains that are responsible for the difference in robustness of STAT3 activation.

### The greater phosphorylation of STAT3 induced by SarA is not due to a lack of SOCS3 binding

Differences between the GBS domains of gp130 and SarA provided an opportunity to determine the molecular mechanism responsible for the supraphysiological STAT3 activation by SarA. First, we looked at the role of negative regulators SOCS3 and SHP-2 (encoded by the gene *PTPN11*) on SarA activation of STAT3. Within the GBS domain of gp130 there is a YxxV SHP-2/SOCS3 binding motif. SHP-2 and SOCS3 are the major negative regulators of the canonical gp130/STAT3 pathway. When they bind to this motif on gp130, they may prevent STAT3 binding to the receptor or promote STAT3 degradation; the mechanism is still unclear.^16–19^ However, SarA does not have the YxxV motif in its GBS domain, suggesting that neither of these negative regulators can bind and downregulate SarA-induced STAT3 phosphorylation.

To test this hypothesis, we generated two FLAG-SarA constructs predicted to impact SHP-2/SOCS3 binding in opposite ways (**Fig. 2A**). The first construct has the YSTV SHP-2/SOCS3 binding motif from the gp130 GBS domain mutated into the wild-type SarA background (FLAG-SarA^YSTV^). For the second construct, the tyrosine of the YSTV motif in the SarA:gp130 chimeric background was mutated to a phenylalanine, breaking the binding function of the motif as SOCS3 binding requires a phosphotyrosine residue (FLAG-SarA:gp130^Y159F^). We overexpressed these mutant constructs in HeLa cells and measured the ratio of phospho-STAT3 to total STAT3 by western blot.

**Figure 2:**
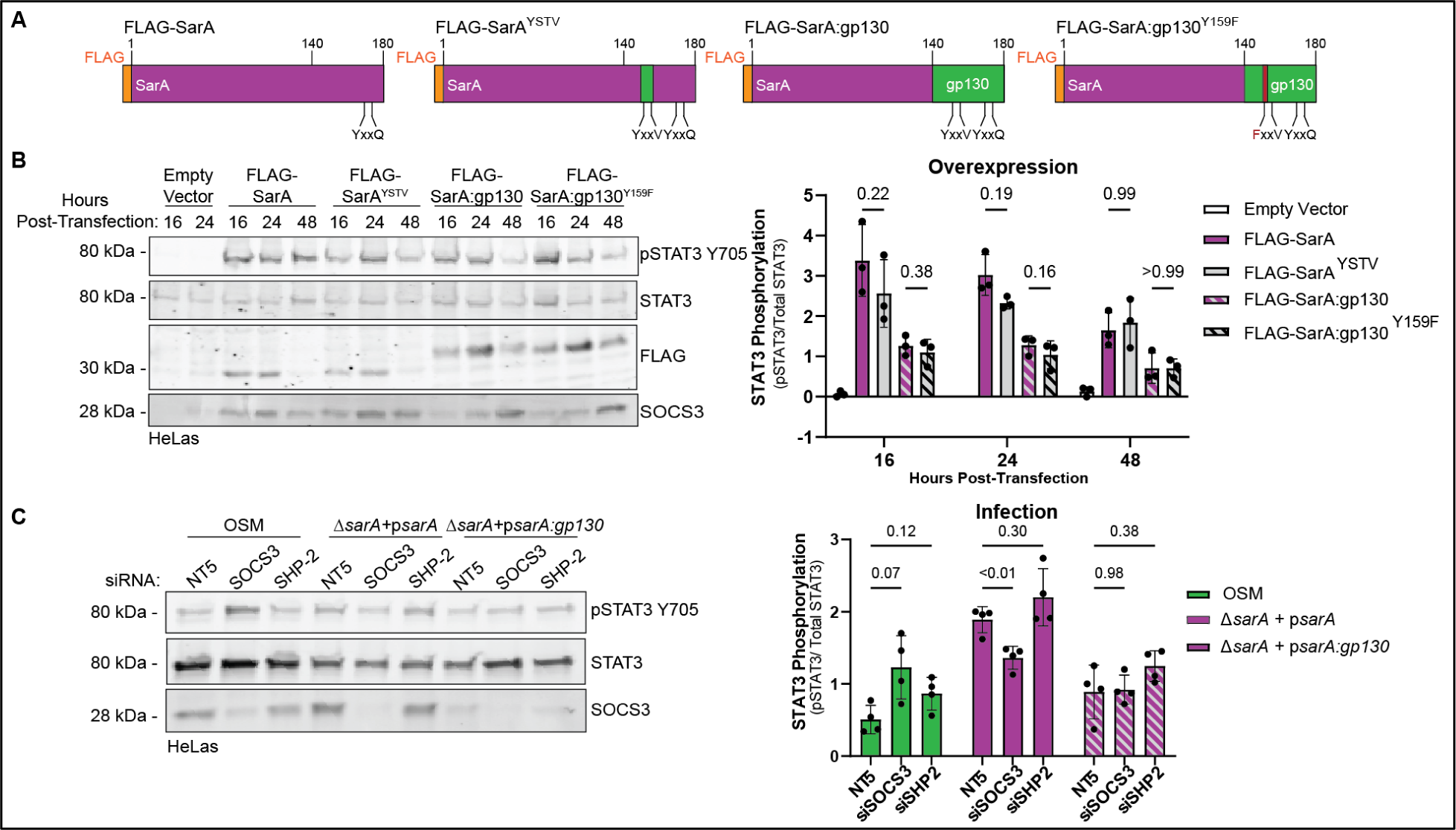
Lack of SOCS3/SHP-2 binding does not explain supraphysiological activation of STAT3 by SarA. (A) Diagrams of FLAG-SarA YxxV mutant constructs used in overexpression experiments. (B) Effects of mutations of SOCS3/SHP-2 binding motifs on phospho-STAT3. Constructs in (A) were overexpressed in HeLa cells. Mutating the YxxV SOCS3/SHP-2 binding motif had no significant impact on STAT3 phosphorylation as measured by western blot. Data are from three experiments. P-values obtained from a 2-way ANOVA with a Tukey’s multiple comparisons test. (C) Knockdown of SOCS3 and SHP-2 do not significantly increase SarA-induced STAT3 phosphorylation. HeLa cells were treated with either a non-targeting control (siGenome non-targeting #5), siSOCS3, or siSHP-2 48 hrs before infection with either a wild-type complemented or p*sarA:gp130* complemented strain of *S.* Typhimurium. STAT3 phosphorylation measured by western blot shows that knocking down negative regulators SOCS3 and SHP-2 were not able to restore p*sarA:gp130* induced activation to wild-type levels. Data are from four experiments. P-values obtained from a 2-way ANOVA with a Tukey’s multiple comparisons test.

Overexpression of the FLAG-SarA^YSTV^ construct did not significantly decrease phospho-STAT3 levels compared to wild-type SarA. Similarly, overexpression of FLAG-SarA:gp130^Y159F^ did not lead to significantly increased phospho-STAT3 protein levels compared to FLAG-SarA:gp130 (**Fig. 2B**). These results demonstrate that altering the SHP-2/SOCS3 binding motif does not significantly impact the function of the GBS.

To confirm these overexpression-based findings, we performed a complementary experiment measuring the effects of RNAi knockdown of SOCS3 and SHP-2 in HeLa cells. Following knockdown of either SOCS3 or SHP-2, cells were treated with oncostatin M (OSM), an IL-6 family cytokine that signals through gp130, or infected with either a p*sarA* or p*sarA:gp130* strain of *S.* Typhimurium for 24 hrs. As predicted, OSM-dependent STAT3 phosphorylation was increased when SOCS3 was depleted (**Fig. 2C**). In contrast, the results show that knocking down SOCS3 or SHP-2 did not increase phospho-STAT3 levels during either p*sarA* or p*sarA:gp130* infection. In fact, through an unknown mechanism, there was a moderate but significant *decrease* in phospho-STAT3 with *SOCS3* knockdown during p*sarA* infection (**Fig. 2C**). Regardless, both the overexpression and loss-of-function infection experiments indicate that the loss of SOCS/SHP2 binding cannot explain the SarA GBS domain’s increased induction of STAT3 phosphorylation compared to the gp130 GBS domain.

### A single amino acid difference controls the robustness of SarA-STAT3 activation

Previously, we determined that the phosphorylated YxxQ STAT3 binding motif in SarA binds to STAT3 approximately 16 times stronger than the phosphorylated motif in gp130 ^6^. This led us to a second hypothesis for how SarA could lead to greater activation of STAT3: amino acid differences near the YxxQ motif might facilitate greater binding affinity, leading to greater phosphorylation of STAT3.

Using homology modeling of STAT3 bound to either SarA or gp130 5mer peptides, we predicted which amino acids in the YxxQ STAT3 binding motif might have the greatest effect on binding affinity. The arginine in the pY+1 position of gp130 (R768) appeared to sterically clash with STAT3 at S54. In contrast the isoleucine in the pY+1 position of SarA (I168) was modeled to form a hydrogen bond with this same S54 residue. Therefore, we generated six FLAG-tagged SarA constructs with mutations in the YxxQ motif (**Fig. 3A**) and overexpressed these constructs in HeLa cells for 24 hours. Mutating the pY+1 position of wild-type SarA from isoleucine to arginine (I168R) significantly decreased phospho-STAT3 levels—down to the level of SarA:gp130. In contrast, mutating the pY+1 position in SarA:gp130 from arginine to isoleucine (R168I) significantly increased phospho-STAT3 to wild-type levels (**Fig. 3B**). Notably, switching the pY+2 position or making a conservative pY+1 change (isoleucine to leucine) had minimal effect on phospho-STAT3 levels.

**Figure 3:**
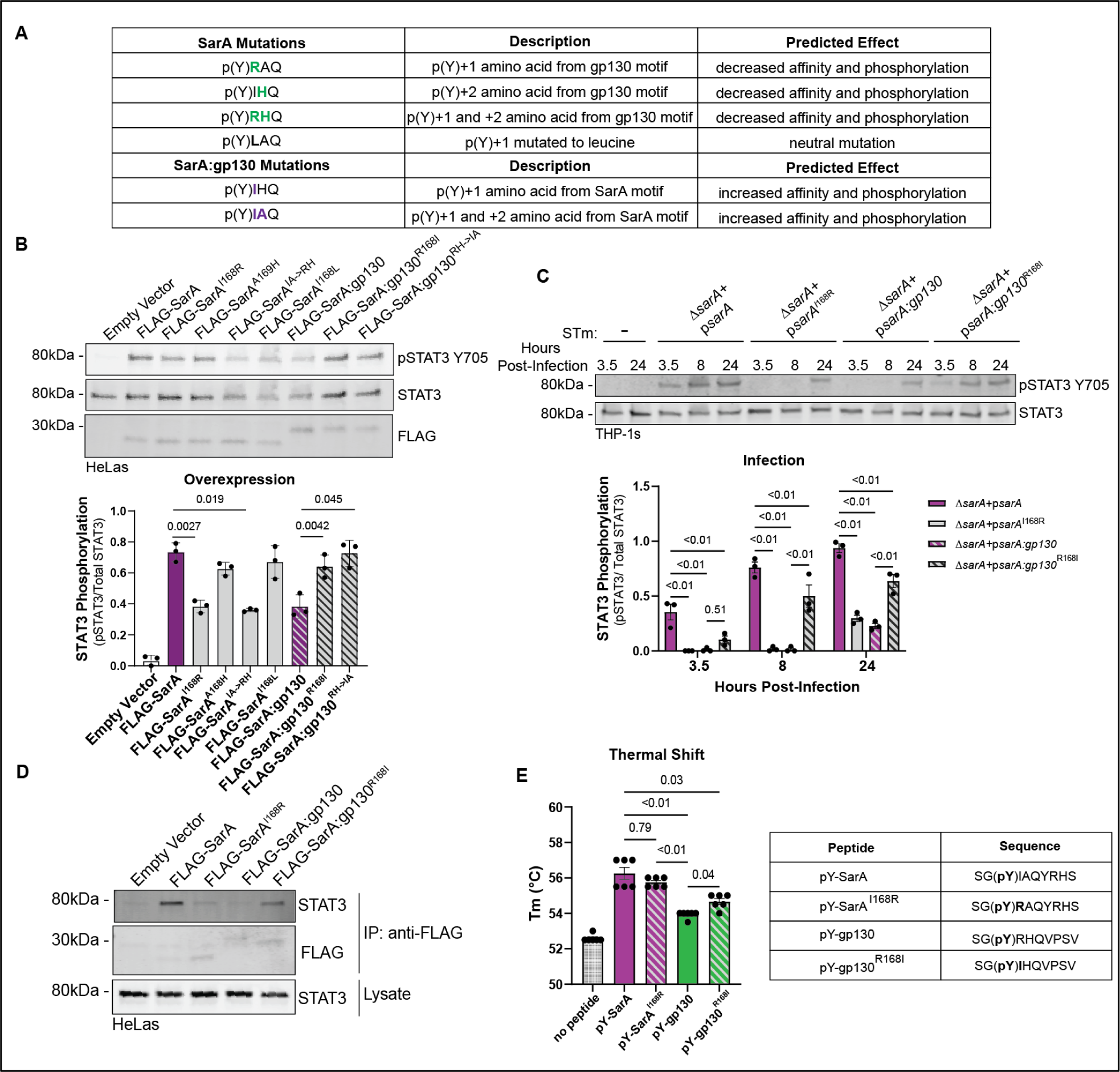
Isoleucine at pY+1 position leads to increased STAT3 binding and phosphorylation. (A) Table of mutations made to YxxQ STAT3 binding motif in FLAG-SarA and FLAG-SarA:gp130 constructs. (B) Effects of mutants on STAT3 phosphorylation. Constructs in (A) were overexpressed in HeLa cells. STAT3 phosphorylation measured by western blot shows that mutating the pY+1 position in SarA from isoleucine to arginine leads to a significant decrease in STAT3 activation. Mutating the same position in SarA:gp130 from arginine to isoleucine restores STAT3 phosphorylation to wild type levels. Mutating the pY+2 position has no significant effect. Data are from three experiments. P-values are from a one-way ANOVA with Dunnett’s multiple comparisons tests comparing FLAG-SarA to its derived mutants and FLAG-SarA:gp130 to its derived mutants. (C) Isoleucine at pY+1 position leads to greater STAT3 phosphorylation during infection. THP-1 cells were infected with wild-type, p*sarA*^I168R^, p*sarA:gp130*, or p*sarA:gp130*^R168I^ complemented *S.* Typhimurium. STAT3 phosphorylation was measured by western blot. Data are from three experiments. P-values are from a 2-way ANOVA with Tukey’s multiple comparisons test. (D) FLAG-SarA constructs with arginine at pY+1 position bind less STAT3. FLAG-SarA, FLAG-SarA^I168R^, FLAG-SarA:gp130, and FLAG-SarA:gp130^R168I^ were overexpressed in HeLa cells, followed by co-immunoprecipitation and probing for bound STAT3 via western blot. Data are representative of three experiments. (E) Mutating the pY+1 position has a minimal effect on STAT3 binding. Purified STAT3 was incubated with 25µM phosphopeptides and a thermal shift assay was used to calculate the melting temperature of the bound peptides as a proxy for binding affinity. Mutating the pY+1 position did not significantly affect melting temperature. Data are from two experiments. P-values are from a Brown-Forsythe and Welch ANOVA tests with Dunnett’s T3 multiple comparisons test.

To confirm this result during *S.* Typhimurium infection, we generated strains of *S.* Typhimurium that express SarA with the same YxxQ mutations. We infected THP-1 cells with these strains and measured phospho-STAT3 levels over the course of a 24-hour infection. Our results show that the p*sarA*^I168R^ mutant strain has significantly slower and lower overall amount of STAT3 phosphorylation during infection compared to the wild type complemented strain; this phenocopies the pattern of phosphorylation during infection with the p*sarA:gp130* expressing strain (**Fig. 3C**). During infection with the p*sarA:gp130*^R168I^ mutant strain, STAT3 phosphorylation was significantly increased compared to infection with the p*sarA:gp130* strain (**Fig. 3C**).

To determine whether the pY+1 mutations affect STAT3 phosphorylation due to changes in STAT3 binding, we performed co-immunoprecipitation experiments using our FLAG-tagged SarA overexpression constructs. The SarA^I168R^ mutant exhibited reduced binding to STAT3, down to the level seen with the SarA:gp130 chimera (**Fig. 3D**). In contrast, the SarA:gp13^R168I^ mutant had levels of STAT3 binding that were similar to the wild-type SarA construct (**Fig. 3D**).

We then tested whether the pY+1 mutations changed the binding affinity to STAT3 as the differences in phosphorylation levels and binding of STAT3 in cells would suggest. Previously, thermal shift assays were used to demonstrate binding of gp130 peptide to STAT3.^20^ Therefore, we developed a thermal shift assay (TSA) to measure the thermostability of recombinant, purified STAT3 in the presence of phosphopeptides derived from the SarA and gp130 STAT3 binding sites. The TSA served as a proxy for direct measurement of binding affinity. The TSA recapitulated the greater affinity of STAT3 for the phospho-SarA peptide compared to the phospho-gp130 peptide. Surprisingly, mutating the peptides at the pY+1 position only resulted in partial and for phospho-SarA, a non-significant effect (**Fig. 3E**), indicating that enhanced binding affinity was not the primary mechanism by which the I168R and R168I mutations were exerting their effect on STAT3 activation.

### Isoleucine at the pY+1 position makes SarA a better substrate for GSK-3 tyrosine phosphorylation

As binding of SarA to STAT3 requires phosphorylation of the tyrosine in the YxxQ motif ^6^, we hypothesized that the reduced binding of STAT3 could be secondary to the level of SarA phosphorylation. Immunoprecipitation of FLAG-SarA demonstrated much greater tyrosine phosphorylation from the constructs with isoleucine in the 168 position (**Fig. 4A**). Therefore, the isoleucine at the pY+1 position in SarA increased the level of SarA phosphorylation to increase STAT3 binding, resulting in levels of activation higher than with gp130.

**Figure 4:**
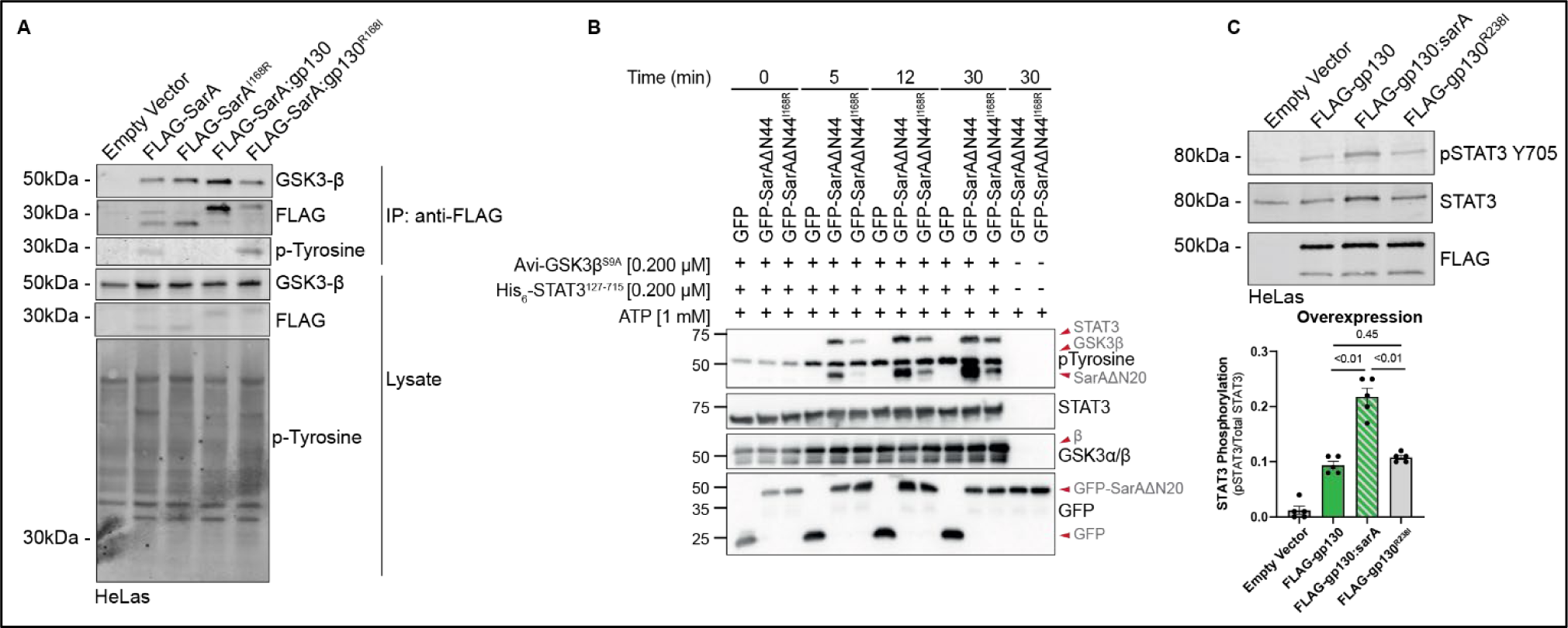
Isoleucine at the pY+1 position makes SarA a better substrate for GSK-3 but has no effect on gp130. (A) Isoleucine at the pY+1 position is required for SarA phosphorylation. Empty vector and panel of FLAG-SarA constructs were overexpressed in HeLa cells and co-immunoprecipitation confirms that while GSK-3 is bound to all constructs, tyrosine phosphorylation of SarA^I168R^ and SarA:gp130 was not detected. Data are representative of two experiments. (B) Mutation of isoleucine at the pY+1 position greatly reduces phosphorylation of SarA by GSK-3. GFP, GFP-SarAΔN44, and GFP-SarAΔN44^I168R^ were expressed in *GSK-3α/β*^-/-^ 293ET cells, immunoprecipitated, and assessed for their ability to be tyrosine phosphorylated by GSK-3β in an in vitro kinase assay containing 1 mM ATP with or without recombinant Avi-GSK-3βS9A (0.2 μM) and His6-STAT3127-715 (0.2 μM). Data are representative of three experiments. (C) Isoleucine at the pY+1 position is not sufficient to induce greater levels of gp130-mediated STAT3 phosphorylation. Empty vector, FLAG-gp130, FLAG-gp130:sarA, and FLAG-gp130^R238I^ were overexpressed in HeLa cells for 24hrs, STAT3 phosphorylation assessed by western blot shows that the R238I single point mutation does not increase pSTAT3 levels compared to wild type gp130. Data are from four experiments. P-values obtained from a Brown-Forsythe and Welch ANOVA with Dunnett’s T3 multiple comparisons test.

This increased phosphorylation suggests that isoleucine at the pY+1 position makes the YxxQ motif a better substrate for GSK-3 tyrosine phosphorylation. While GSK-3 is canonically a serine-threonine kinase, it does auto-phosphorylate its activation loop at Y216 in GSK-3β.^21^ Previously, the Thurston lab demonstrated the direct phosphorylation of SarA by GSK-3, revealing the first exogenous substrate of GSK-3 tyrosine phosphorylation.^7^ Therefore, we conducted an *in vitro* kinase assay using SarA containing isoleucine or arginine at the pY+1 position as well as recombinant STAT3 and GSK-3β. We observed greater tyrosine phosphorylation of SarA by GSK-3 with isoleucine present at the pY+1 position at all time points tested, and this corresponded with greater tyrosine phosphorylation of STAT3(**Fig. 4B**). Thus, isoleucine at pY+1 renders SarA a better substrate for GSK-3 tyrosine phosphorylation.

### Constitutively active gp130 does not induce greater STAT3 activation with isoleucine at the pY+1 position

The altered levels of STAT3 phosphorylation induced with the chimeric SarA constructs depends on the presence of GSK-3 as the cognate kinase. Therefore, it was unclear whether isoleucine at the pY+1 position would have a similar effect within gp130. To test this, we made an R238I mutation in the constitutively active gp130 dimer construct. While replacing the entire 39-amino acid GBS segment with the homologous SarA sequence resulted in greater phospho-STAT3 levels (consistent with our previous studies ^6^, overexpression of the R238I mutant in HeLa cells did not alter phopho-STAT3 levels compared to the wild-type gp130 construct (**Fig. 4C**).

### Tuning SarA activation of STAT3 controls the bias of target genes

Does supraphysiological activation of STAT3 impact downstream transcriptional targets? To test this, we infected THP-1s with the YxxQ mutant strains of *S.* Typhimurium and measured the transcription and production of downstream anti-inflammatory STAT3 targets. Our results demonstrate that the p*sarA*^I168R^ mutation leads to reduced transcription of anti-inflammatory genes down to p*sarA:gp130* levels, and increased transcription is restored with the p*sarA:gp130*^R168I^ mutation (**Fig. 5A**). Infection with the p*sarA*^I168R^ mutant strain also leads to significantly lower production of IL-10 compared to wild-type, whereas infection with the p*sarA:gp130*^R168I^ mutant leads to significantly increased IL-10 compared to the p*sarA:gp130* strain (**Fig. 5B**).

**Figure 5:**
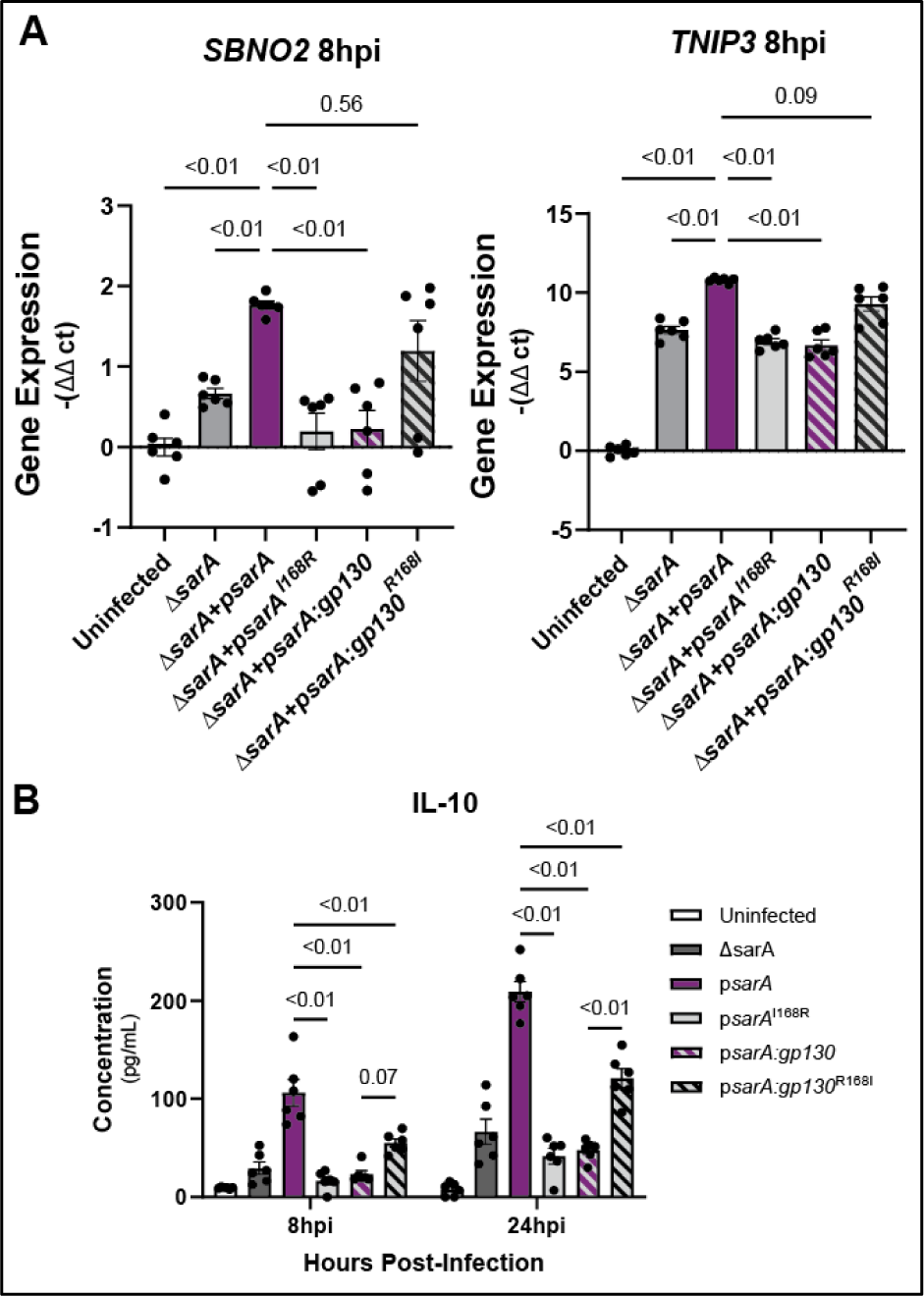
Isoleucine at pY+1 position promotes anti-inflammatory signaling bias. (A) Arginine in the pY+1 position leads to decreased expression of anti-inflammatory genes during infection and isoleucine at this position rescues expression of these genes. THP-1 cells were infected with Δ*sarA*, wild-type complemented, p*sarA*^I168R^, p*sarA:gp130*, or p*sarA:gp130*^R168I^ *S.* Typhimurium. RNA was collected from cells 8hrs post-infection and expression levels of *SBNO2* and *TNIP3* were measured via qPCR. Data are from three experiments. P-values obtained from a Brown-Forsythe and Welch ANOVA with Dunnett’s T3 multiple comparisons test. (B) Arginine in the pY+1 position leads to decreased production of IL-10 during infection and is rescued with isoleucine at the position. THP-1 cells were infected with Δ*sarA*, wild-type complemented, p*sarA*^I168R^, p*sarA:gp130*, or p*sarA:gp130*^R168I^ *S.* Typhimurium. Cell supernatant was collected at 8 hrs and 24 hrs post-infection and IL-10 was measured via ELISA. Data are from three experiments. P-values obtained from a two-way ANOVA with a Sidak’s multiple comparison test.

### Evolution of STAT3 activation by SarA vs. gp130

Our results show that the amino acid in the pY+1 position of the YxxQ motif in SarA leads to increased phosphorylation by GSK-3, which causes increased STAT3 binding and increased phospho-STAT3 levels compared to the residue in the same position in gp130. SarA is present on the Gifsy-1 prophage ^5^ and can therefore be acquired through horizontal gene transfer among serovars. BLASTp of SarA in over 20,000 *Salmonella* strains ^22^, demonstrated the presence of SarA in 4,554 strains, including members of the Arizonae and Houtenae subspecies that primarily infect birds and reptiles (**Fig. 6A**). In these 4,554 isolates, the pY+1 position is invariably isoleucine (**Fig. 6B**). Thus, all examined *Salmonella* strains that encode for SarA have retained the isoleucine at the pY+1 position that makes SarA’s YxxQ motif a better substrate for GSK-3, facilitating supraphysiological activation of STAT3.

**Figure 6:**
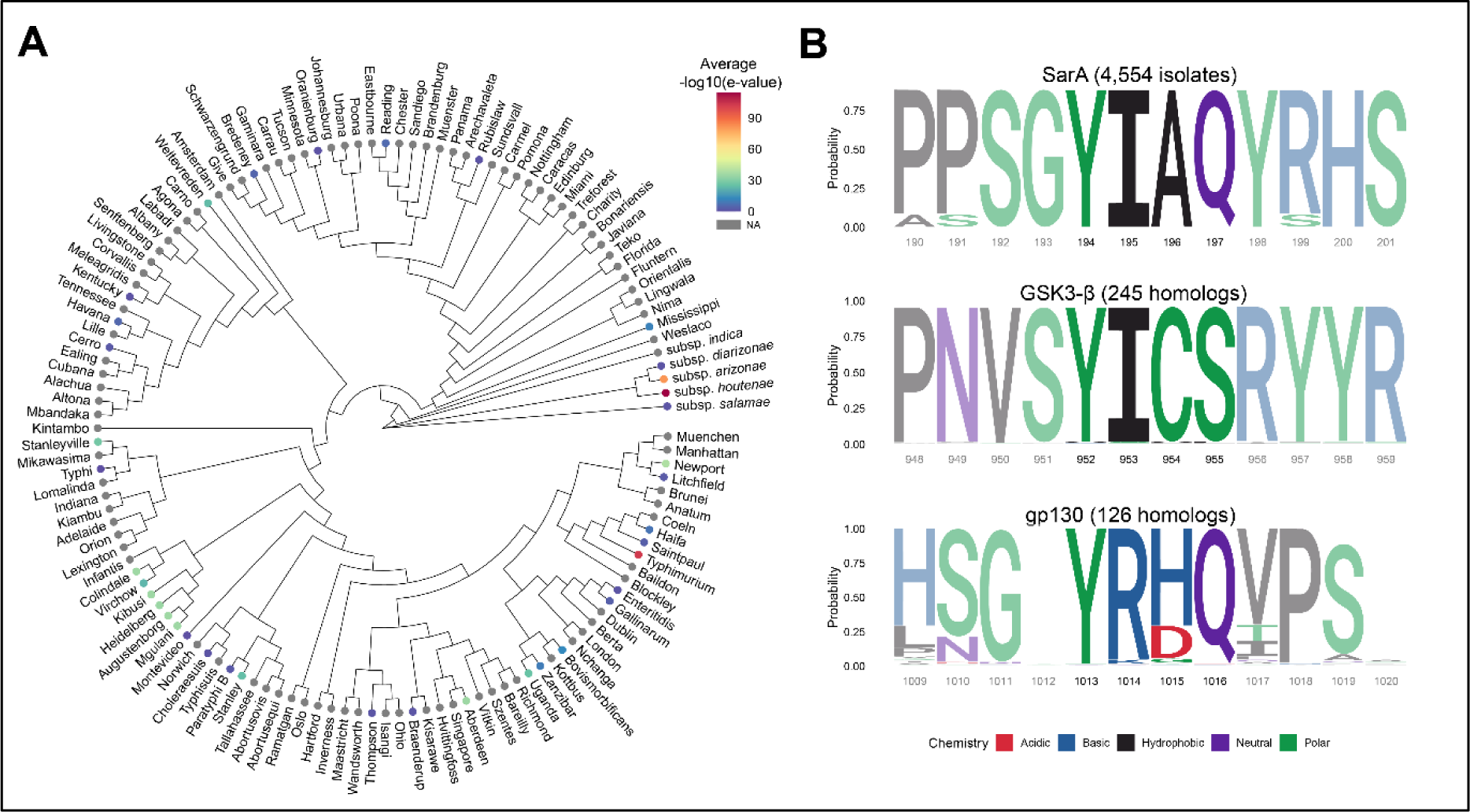
SarA is variably present in *Salmonella* serovars but isoleucine at pY+1 position is always conserved. (A) SarA is present in diverse *S. enterica* subspecies and serovars. Mean E-values (indicated by color) from BLASTp of SarA in 21,223 *Salmonella* isolates were merged with an existing whole genome maximum-likelihood phylogeny (Worley et al., 2018). Serovars without corresponding BLASTp data are in gray. (B) Isoleucine at pY+1 is invariable across *Salmonella* isolates that contain SarA and homologs of the human protein GSK-3β. Sequence logos were derived from multiple sequence alignment and reflect the relative frequency of each residue across all aligned sequences. The YxxQ motif that is homologous between SarA and IL6ST is highlighted, as well as the known autophosphorylation site in the kinase GSK-3β. Known chemical classification of each residue is indicated by color.

Isoleucine at this pY+1 position also shows conservation at the autophosphorylation site of GSK-3β across 244 homologs of human GSK-3β across 197 species extending back to yeast (**Fig. 6B, Table S4**). Thus, maintaining high activity of this autophosphorylation event appears to be under evolutionary constraint, consistent with GSK-3’s high baseline activity in unstimulated cells.^23^ In contrast, multiple sequence alignment comparison of 133 homologs of human gp130 showed the pY+1 position is almost invariably arginine across 131 species extending back to fish (**Fig. 6B**). The exceptions are changes to the similarly basic lysine residue in spotter gar, elephant shark, Old Calabar mormyrid elephantfish, and Asian bonytongue fish. This indicates conservation at the pY+1 position extending to the common ancestor of fish and tetrapods in the Devonian period ∼400 million years ago.^24^ While R238I is not sufficient to increase the activity of gp130 (see **Fig. 4C**), greater activation of STAT3 by gp130 is possible with further mutagenesis of the GBS based on the gp130-SarA chimeric construct. Therefore, gp130 appears to have evolved to mediate an intermediate level of activation of STAT3.

## Discussion

Many intracellular pathogens, including both bacteria and viruses, have been shown to activate anti-inflammatory pathways during infection to evade host pro-inflammatory attacks.^25,26^ In this paper, we demonstrate that a single amino acid residue in the YxxQ motif of *S.* Typhimurium effector SarA controls the level of STAT3 activation to induce IL-10 production and cause an anti-inflammatory signaling bias in infected cells. SarA signaling through STAT3 drives M2 macrophage polarization during systemic *Salmonella* Typhimurium infection,^7–9^ while IL-10 secreted by T and B cells is required for dissemination of *S.* Typhimurium from the gut to systemic sites of infection in a mouse model.^27^ SarA-induced IL-10 production could lead to non-cell autonomous responses that benefit the bacteria, such as decreasing production of pro-inflammatory cytokines by bystander cells, preventing the recruitment of other immune cell populations, and reducing T-cell responses. The relative contributions of these possible effects of SarA during *Salmonella* infection is an important direction for future work.

Beyond *Salmonella,* other bacteria also hijack STAT3 through different mechanisms. *Bartonella hensalae*, which causes cat scratch disease in humans, employs the effector BepD to activate STAT3 in a c-Abl kinase-dependent mechanism to suppress TNFα signaling and stimulate IL-10 production.^28^ While BepB signaling in this context has an anti-inflammatory bias similar to SarA activation, it uses a kinase that, unlike GSK-3, is not active at baseline.^29^ *Helicobacter pylori*, which typically causes chronic infection of the stomach, also manipulates STAT3 signaling. Its effector CagA is notable for being able to either activate *or* suppress IL-6/gp130 signaling depending on the effector’s phosphorylation state.^30^ CagA has multiple phosphorylation sites that have been shown to be phosphorylated by c-Src and c-Abl kinase families,^31^ suggesting that the effector’s signaling pathway can be tightly controlled and different combinations of phosphorylation may occur in dynamic *H. pylori* populations during infection. Further, as CagA acts through IL-6 and gp130 signaling for STAT3 activation, the features of this host signaling pathway that reduce STAT3 signaling (JAK-dependence, inhibition by SOCS proteins) are still at play in this context. Thus, while multiple pathogens can activate STAT3, the magnitude of STAT3 activation may be adapted to each particular pathogen. Whereas supraphysiological activation of STAT3 with strong anti-inflammatory bias may be adaptive for SarA-containing *Salmonella* serovars that typically cause acute gastroenteritis in human hosts, more nuanced control may be utilized by *H. pylori* in causing chronic infection. The variable presence of SarA among *Salmonella* serovars suggests complex bacterial adaptation to different host species and requires further investigation into how triggering supraphysiological activation of STAT3 affects various pathologies.

Effector activation of STAT3 may at first seem counterintuitive, as the host is already producing IL-6 in response to *Salmonella* infection,^32–34^ and we previously observed activation of STAT3 above uninfected levels even in the absence of SarA ^5^. SarA appears to exploit a homeostatic mechanism, whereby the host uses the same transcription factor (STAT3) to induce pro-inflammatory, and at greater levels of activation, anti-inflammatory targets. Short, rapid phosphorylation of STAT3 via IL-6 signaling leads to pro-inflammatory target genes being activated. In contrast, continued and more robust activation of this same transcription factor via IL-10 signaling leads to activation of anti-inflammatory target genes.^14^ Understanding how a limited set of shared proteins mediate the very different effects of IL6, IL-10, and other cytokines that induce JAK/STAT signaling is still a very active area of research.^35^ One mechanism underlying this difference between IL-6 and IL-10 signaling is responsiveness to negative regulators: SHP-2 and SOCS3 bind to gp130 to inhibit signaling, but neither of these proteins can downregulate IL-10 signaling.^36^ It has also been shown that deletion of SOCS3 or mutation of its binding site in gp130 results in IL-6 displaying an anti-inflammatory bias similar to IL-10 signaling.^11,18,37^

We initially hypothesized that SarA was able to manipulate STAT3 phosphorylation dynamics by avoiding these negative regulators, similar to IL-10. Instead, we found that *Salmonella* mediated activation of STAT3, by co-opting the constitutively active kinase GSK-3 is what determines the downstream signaling bias. Both IL-6 and IL-10 signaling utilize strict control of the JAK kinases that activate STAT3, forming an active complex with JAK and STAT3 upon cytokine stimulation. GSK-3 is a serine/threonine kinase with over a hundred known substrates and plays an important role in metabolic regulation, proliferation, and inflammation.^38,39^ Despite exhibiting strict serine/threonine activity in its mature state, GSK-3 is sometimes classed as a dual specificity kinase, as it auto-tyrosine phosphorylates on a tyrosine residue in its activation loop during translation.^21^ Prior to the discovery of SarA and STAT3 phosphorylation by GSK-3,^7^ autophosphorylation at the YICS sequence in GSK-3’s activation loop – which is required for full protein activity – was the only known substrate for this activity.^21^ Our results show that SarA has evolved to match the GSK-3 autophosphorylation site to facilitate constitutive STAT3 activation in a cytokine-independent manner. Notably, the phosphorylation site in STAT3 targeted by GSK-3 has leucine at the pY+1 position (L706), a conservative change, that at least in the SarA YxxQ motif, has no effect on phosphorylation (see Figure 3B). The autophosphorylation of GSK-3 occurs primarily during the folding process, in a chaperone-dependent manner.^21^ How SarA is able to re-awaken this activity is unknown, but in addition to having the optimal substrate sequence with isoleucine in the pY+1 position, there may be structural contributors to this. The GBS region in SarA is rich in prolines and serines (12 of 39 residues) which we speculate may result in an extended, poorly structured sequence. While our work reveals a molecular basis as to how SarA is able to activate STAT3 to supraphysiological levels, future work will be needed to define the structural basis that underlies these findings and whether there are more substrates for the tyrosine kinase activity of GSK-3.

## Methods

### Plasmids

Plasmids are listed in Table S1. The pcDNA3.1-FLAG*-sarA*, pcDNA3.1-FLAG*-sarA:gp130* chimera, pcDNA3.1-FLAG-*gp130dimer*, pcDNA3.1-FLAG*-gp130:sarA* chimera codon-optimized for mammalian expression were previously described in and available from Addgene. Site-directed mutagenesis was carried out using Quikchange Lightning Site-Directed Mutagenesis Kit (Agilent, # 210518) (primers in Table S2). Note that amino acid numbering used throughout this manuscript is based on the Uniprot entry for SarA (A0A0F6B506). Panagi et al. noted that the translational start very likely occurs 72 nucleotides (24 amino acid residues) later than indicated ^7^, but as all of our human codon-optimized SarA plasmids contain these additional bases, we have indicated the critical isoleucine residue as being at amino acid position 168.

### Mammalian and bacterial cell culture

THP-1s were cultured at 37°C in 5% CO2 in RPMI 1640 media (Invitrogen) supplemented with 10% heat-inactivated FBS (Thermo-Fisher), 2 μM glutamine, 100 U/mL penicillin-G, and 100 mg/mL streptomycin. HeLa cells (Duke Cell Culture Facility) were grown in high glucose DMEM media supplemented with 10% FBS, 1mM glutamine, 100 U/mL penicillin-G, and 100mg/mL streptomycin. Cells used for *Salmonella* gentamicin protection assays were grown without antibiotics for at least one hour prior to infection.

All *Salmonella* strains are derived from the *S*. Typhimurium strain 14028s and are listed in Table S3, and all plasmids are listed in Table S1. For infection of cells or mice, bacteria were grown overnight in LB broth (Miller formulation, BD), subcultured 1:33 in 1mL cultures, and grown for an additional 160 minutes at 37°C shaking at 250 RPM. Ampicillin was added to LB at 100 μg/mL, kanamycin at 50 μg/mL, chloramphenicol at 20 μg/mL.

### Transfections

HeLa cells were seeded at 1.75 × 10^5^ cells per well in 6-well TC-treated plates or 7.5 × 10^4^ cells in 24-well TC-treated plates 24 hours before transfection. Transfection was accomplished with Lipofectamine 3000 (Thermo Fisher, Catalog #L3000008). Cells were harvested 16, 24, and 48 hours post transfection.

### Immunoprecipitation

HeLa cells were seeded at 1.75 × 10^6^ cells per 10 cm dish 24 hours before transfection with codon-optimized pcDNA3.1-Flag-sarA, pcDNA3.1-FLAG-sarA:gp130, pcDNA3.1-FLAG-sarA^I168R^, or pcDNA3.1-FLAG-sarA:gp130^R168I^ as described above. Four plates per condition were lysed in 1mL of 0.1% Triton X-100, 50mM Tris pH 7.4, 150mM NaCl, cOmplete Mini protease inhibitor cocktail (Millipore Sigma #11836170001), 10mM NaF, and 1-mM Na Orthovanadate. Lysate was incubated with anti-FLAG M2 magnetic beads (Millipore Sigma #M8823) for 4 hrs while rotating at 4C then washed 3x with 200µL of lysis buffer using a DynaMag-2 magnet (Invitrogen #12321D). Bound protein was eluted with 45µL of 0.1-M Glycine-HCl buffer pH 3 and neutralized with 5µL of 0.5M Tris-HCl pH 7.4 1.5M NaCl. Eluted proteins were analyzed by immunoblot.

### Immunoblotting

Cells were lysed with RIPA lysis buffer: 50mM Tris-HCl pH 7.4, 150mM NaCl, 0.1% SDS, 0.5% NaDeoxycholate, 1% Triton X-100, cOmplete Mini protease inhibitor cocktail (Millipore Sigma #11836170001) 10mM NaF, and 1-mM Na orthovanadate. SDS-PAGE was performed using Mini-PROTEAN TGX Stain-Free Precast 4%–20% gels (Bio-Rad #456-8096). The gels’ stain-free dye was activated by a 5-min UV exposure and protein was transferred to Immun-Blot low-fluorescence PVDF membrane (Bio-Rad #162-0264) using Hoefer TE77X. Primary antibodies used were: anti-FLAG M2 (Sigma F3165, RRID:AB_259529), anti-pY705-STAT3 clone D3A7 (CST #9145, RRID:AB_2491009), anti-STAT3 clone 124H6 (CST #9139, RRID:AB_331757) and anti-SOCS3 polyclonal (proteintech #14025-1-AP, RRID: AB_10597854). Blots were developed using LI-COR infrared secondary antibodies (IRDye 800CW Donkey anti-rabbit IgG and IRDye 680LT Donkey anti-mouse IgG) and imaged on a LI-COR Odyssey Classic. Total protein was measured following a 30-sec UV exposure. Band integrated intensity was quantified using Image Studio^TM^ v5.5. Background was subtracted using the median of the top and bottom of the band. STAT3 phosphorylation was calculated as pSTAT3-Y705 integrated intensity over total STAT3 integrated intensity and then set relative to a control as specified in the figure.

### Infection assays

Assaying infection of THP-1s and HeLa cells was done as previously described ^40^. Overnight cultures of *Salmonella* in LB media were subcultured 1:33 and grown for 160 minutes at 37°C and 250 rpm. THP-1s were seeded at either 100,000 in 100 μl in 96-well or 500,000 in 500ul in 24-well non-TC plates and infected at <u>multiplicity of infection</u> (MOI) 10. HeLa cells at 300,000 in 2 mL were infected at MOI 5 in 6-well TC-treated plates.

Gentamicin was added 1 hpi at 50 μg/mL to kill the extracellular bacteria. For 24-hr incubations, gentamicin was diluted to 15 μg/mL at 2 hpi. To induce GFP, IPTG was added 75 min prior to the desired timepoint. Infection and cell death were measured with a Guava EasyCyte Plus flow cytometer (Millipore). Cell death was measured by 7AAD (7-aminoactinomycin D; Enzo Life Sciences) staining. IL-10 protein in the THP-1 supernatant at 5, 10, 20, 25, and 30hpi was measured by human IL-10 <u>ELISA</u> (R&D systems Catalog #DY217B).

### RNAi experiments

HeLa cells at 37,500 in 24-well TC-treated plates were treated for 24 hours with 5picomoles/well total siRNA from either non-targeting siGENOME (Dharmacon) siRNA #5 (NT5; #D-001210-05) or SMARTpool directed against human *SOCS3* (#M-004299-02-0005) and human *PTPN11* (#M-003947-01-0005). Infections were then conducted 48 hours after initial addition of siRNA as described above.

Simultaneously, knockdown was confirmed in each experiment by qPCR. Briefly, RNA was harvested using a Rneasy kit (Qiagen), cDNA was generated with iScript (Bio-Rad), and qPCR was performed by using iTaq Universal Probes Supermix (Bio-Rad) and a QuantStudio 3 thermo cycler (Applied Biosystems). Primers are listed in Table S2. The cycling conditions were as follows: 50°C for 2 minutes, 95°C for 10 minutes, and 40 cycles of 95°C for 15 seconds followed by 60°C for 1 minute. All qPCR was run in technical duplicate or triplicate. The comparative threshold cycle (CT) was used to quantify transcripts, with the ribosomal 18s gene (RNA18S5) serving as the housekeeping control. ΔCT values were calculated by subtracting the CT value of the control gene from the target gene, and the ΔΔCT was calculated by subtracting the non-targeting siRNA ΔCT from the targeting siRNA ΔCT value. Fold change represents 2-ΔΔCT.

### Thermal shift assay

5 µM STAT3^136–705^ (purified as described in ^6^) was incubated with 25µM phosphopeptides (Genscript, Piscataway, New Jersey) from the binding sites of gp130 (SGpYRHQVPSV), SarA (SGpYIAQYRHS), SarA^I168R^ (SGpYRAQYRHS) and gp130^R168I^ (SGpYIHQVPSV) at room temperature for 5 minutes in an assay buffer containing 10 mM Tris-HCl, pH 8.0; 50 mM NaCl; 10% glycerol; 5 mM β-mercaptoethanol. Water was used as the vehicle control. Subsequently, the STAT3 and peptide mixtures were added to 1X Glomelt^TM^ (Biotium #33021-1, Fremont, California) a dye that binds hydrophobic residues and fluoresces upon protein denaturation due to an increase in heat. The combined assay volume per well was 20 µL. There were 3 replicates per peptide per plate. STAT3 thermal shift assays (TSA) were performed in a 96-well PCR plate (Hard-shell and thin wall PCR plates, Bio-Rad, Hercules, California). Temperature gradient from 25 to 95 °C (increasing ramp rate of 0.05 °C/sec) was completed in a Bio-Rad CFX Connect Real-Time System, following the general guidelines in the Glomelt^TM^ kit protocol. The slope of the first derivative of the fluorescence vs. temperature curve was used to calculate the melting temperature (Tm).

### Kinase Assay

GSK-3α/β-/-293ET cells (Panagi et al., 2020) seeded in a 6-well format were transiently transfected with ptCMV.GFP, ptCMV.GFP-SarAΔΝ44 or ptCMV.GFP-SarAΔΝ20I168R for 48 hrs. The sequence encoding the N-terminal 20 [or 44] amino acid residues of SarA were deleted, as they do not appear to affect activity, and this removes an additional tyrosine phosphorylation site to simplify interpretation. 800 ng of plasmid was transfected in each well of cells and four wells were used per condition. Post-transfection, cells from the same transfection condition were pooled together and lysed in 1.4 mL Lysis Buffer (150 mM NaCl, 0.3% Triton X-100, 20 mM Tris-Cl pH 7.4, 5% Glycerol, 5 mM EDTA), supplemented with a Protease Inhibitor Cocktail (cOmplete™, Roche), and clarified by centrifugation at 17,000 x g for 10 min at 4 °C. Thereafter, GFP-tagged proteins were enriched on beads by GFP-immunoprecipitation. Briefly, GFP-Trap agarose beads (ChromoTek) equilibrated in cold lysis buffer were incubated with the post-nuclear supernatant fraction for 3-5 h at 4°C with rotation. Post-incubation, beads were washed three times in 1 mL Lysis Buffer and twice in 1 mL 1x Kinase Buffer (Cell Signaling) supplemented with Phosphatase Inhibitors (PhosSTOP, Roche). Beads containing GFP-tagged targets were distributed evenly in kinase reaction tubes and resuspended in 50 μL 1x Kinase Buffer containing 1 mM ATP (ThermoFisher) with or without 0.2 μM of non-phosphorylated Avi-GSK-3βS9A and 0.2 μM His6-STAT3127-715 (Panagi et al., 2020). Kinase reactions were carried out at 30°C with agitation at 600 RPM in a PCMT Grant-bio Thermo-Shaker and stopped at the indicated time point by adding 5x SDS loading buffer (312.5 mM Tris-Cl pH 6.8, 10% SDS, 25% Glycerol, Bromophenol Blue) containing 20% β-Mercaptoethanol. The samples were then boiled at 95 °C for 7 min, centrifuged at 1500 x g for 1 min and eluted proteins were subjected to immunoblot analysis.

### Multiple sequence comparison

*Salmonella* genomes deposited in the PubMLST database (n=21,223) were tested for presence of sarA by conducting BLASTp pairwise alignment in PubMLST ^22^. Sequences from samples with a resulting sarA alignment length of >100 residues and a BLAST E-value < 0.001 were aligned using the CLUSTAL W algorithm ^41^ executed through the R package *msa* v.1.34.0 ^42^.

Mean BLASTp E-values from serovars containing samples passing the above criteria were merged with an existing *Salmonella* maximum-likelihood phylogenetic tree described in ^43^ using the R packages *treeio* v.1.26.0 ^44^ and *ggtree* v.3.10.0.^45^

Protein sequences of IL6ST and GSK-3β homologs from Ensembl ^46^ were aligned using the CLUSTAL W algorithm as described above. Sequence logos were generated using the R package *ggseqlogo* v.0.1 ^47^.

### Statistics

Descriptive statistics were performed with GraphPad Prism v10 (GraphPad Software, US) or R v4.3.1 (R Core Team) using tidyverse^48^, magrittr^49^, ggpubr^50^, and venn^51^ packages. The size of each study or number of replicates, along with the statistical tests performed can be found in figure legends. *In vitro* inter-experimental variability was removed prior to data visualization or statistical analysis by making experimental means equal to the grand mean by multiplying all values within each experiment by a normalization constant. These constants were calculated by dividing the mean of all experiments by mean of each specific experiment. Bar graphs represented the mean ± SEM (standard error of mean), unless otherwise noted.

## Acknowledgements

We thank members of the Ko lab for useful discussion. D.C.K., M.R.G., A.G.J. and R.E.L. were supported by NIH R01AI118903. T.L.M.T. was supported by a Biotechnology and Biological Sciences Research Council (BBSRC) David Phillips Fellowship (BB/R011834/1), a Medical Research Council (MRC) Research Grant MR/V031058/1, which also funded J.M., and a European Research Council grant funded by the Engineering and Physical Sciences Research Council EP/X02377X/1 to T.L.M.T., which also funded I.P. E.J.W was supported by the Duke Next Generation Fellowship funded by the Duke Science and Technology Institute. All schematic images were generated using Biorender.com and figures were made with Adobe Illustrator.

## Author Contributions

Conceptualization, M.R.G. and D.C.K.; formal analysis, M.R.G and A.G.J.; investigation, M.R.G., A.G.J., I.P., E.J.W., R.E.L., and D.C.K.; funding acquisition, T.L.M.T. and D.C.K.; supervision, R.G.B., T.L.M.T., and D.C.K.; resources, M.R.G, A.G.J, I.P., R.E.L., J.H.M, E.J.W., R.G.B.,T.L.M.T., and D.C.K.; writing – original draft, M.R.G and D.C.K.; writing – review & editing, M.R.G., A.G.J, R.E.L, I.P., E.J.W., T.L.M.T., and D.C.K.

## Declaration of interests

The authors declare no competing interests.

## Notes

### Competing Interest Statement

The authors have declared no competing interest.

