## Supplemental Figures and Tables for "A single amino acid in the *Salmonella* effector SarA/SteE triggers supraphysiological activation of STAT3 for anti-inflammatory target gene expression"

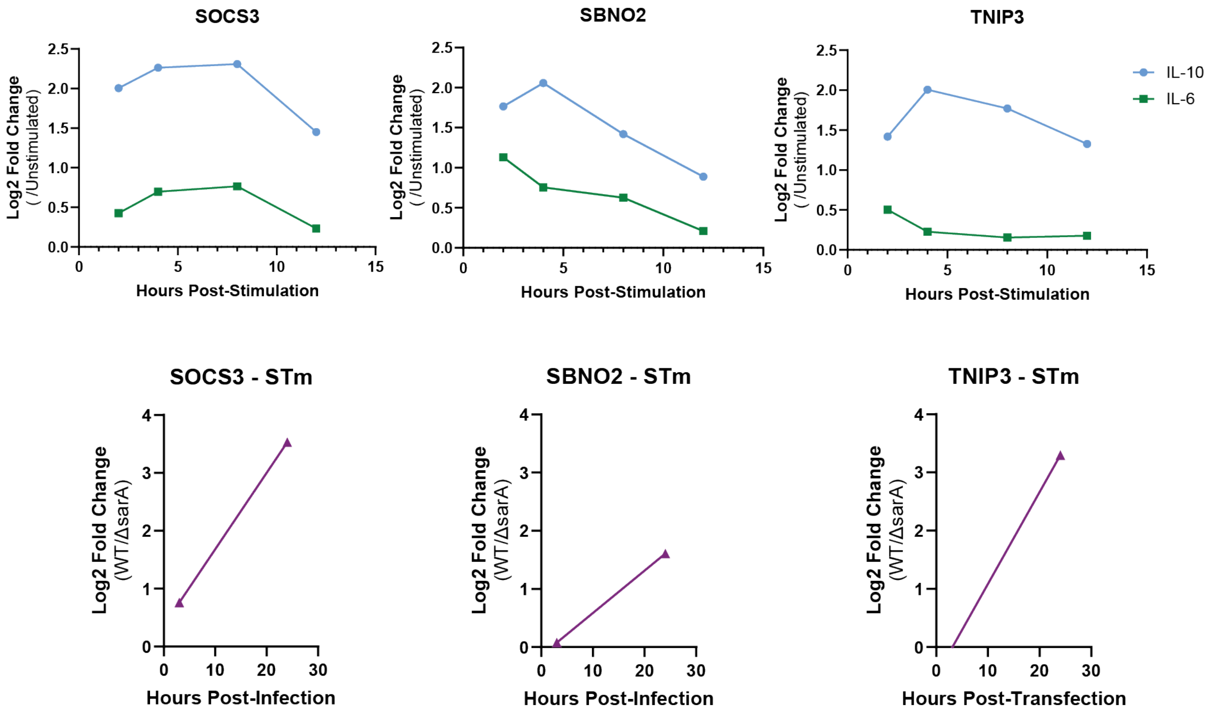

**Supplementary Figure 1: Expression of anti-inflammatory genes after IL10 stimulation, IL-6 stimulation, or *S.* Typhimurium infection.** Log2-fold changes of *SOCS3*, *SBNO2*, and *TNIP3* are from previously published transcriptomic datasets using IL-6 or IL-10 stimulation (Braun et al., 2013) and infection with wild-type or Δ*sarA S.* Typhimurium (Jaslow et al., 2018).

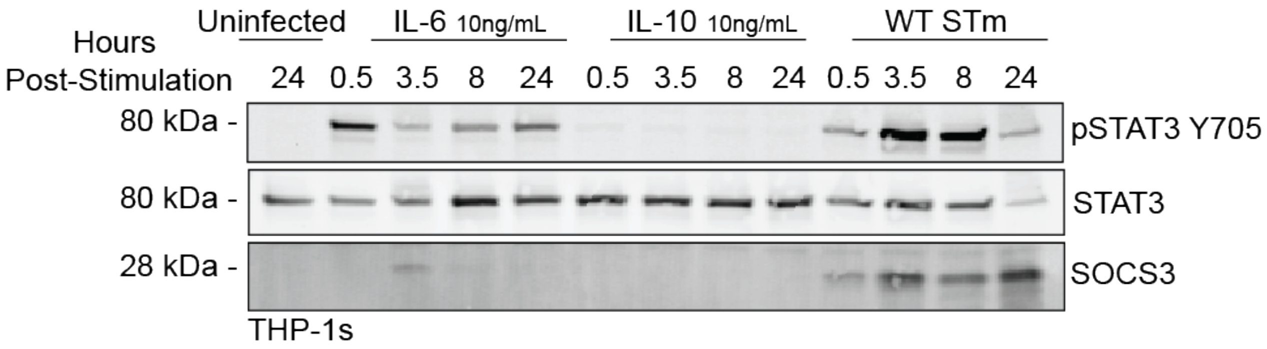

**Figure S2: THP-1 cells have minimal STAT3 phosphorlyation in response to IL-10 stimulation.** THP-1 monocytes were stimulated for the time indicated with IL-6, IL-10, or infected with wild-type *S.* Typhimurium (MOI10).

| Plasmid | Resistance | In E. coli Strain | Notes |
|---|---|---|---|
| pEGFP-C1 | Kan | DCK53 | |
| pcDNA3 | Amp | DCK77 | |
| pcDNA3-FLAG-*sarA* | Amp | DCK796 | |
| pcDNA3-FLAG-*sarA*$^{YSTV}$ | Amp | DCK1150 | |
| pcDNA3-FLAG-*sarA*$^{I168R}$ | Amp | DCK1201 | |
| pcDNA3-FLAG-*sarA*$^{A169H}$ | Amp | DCK1202 | |
| pcDNA3-FLAG-*sarA*$^{IA\text{->}RH}$ | Amp | DCK1203 | |
| pcDNA3-FLAG-*sarA*$^{I169L}$ | Amp | DCK1204 | |
| pcDNA3-FLAG-*sarA:gp130* | Amp | DCK801 | |
| pcDNA3-FLAG-*sarA:gp130*$^{Y159F}$ | Amp | DCK1151 | |
| pcDNA3-FLAG-*sarA:gp130*$^{R168I}$ | Amp | DCK1205 | |
| pcDNA3-FLAG-*sarA:gp130*$^{RH\text{->}IA}$ | Amp | DCK1206 | |
| pcDNA3-FLAG-*gp130dimer* | Amp | DCK824 | |
| pcDNA3-FLAG-*gp130dimer*$^{R\text{->}I}$ | Amp | DCK1254 | |
| pcDNA3-FLAG-*gp130dimer:sarA* | Amp | DCK825 | |
| pWSK129 | Kan | DCK827 | |
| pWSK129-*sarA* | Kan | DCK809 | |
| pWSK129-*sarA*$^{I168R}$ | Kan | DCK1221 | |
| pWSK129-*sarA:gp130* | Kan | DCK852 | |
| pWSK129-*sarA:gp130*$^{R168I}$ | Kan | DCK1220 | |
| ptCMV.GFP | | | From Thurston Lab, Panagi et al., 2020 |
| ptCMV.GFP-*SarAΔ20* | | | From Thurston Lab, Panagi et al., 2020 |
| ptCMV.GFP-*SarAΔ20*$^{I168R}$ | | | From Thurston Lab |

#### Table S1: Plasmids

| Designation | Name | Purpose | Sequence |
|---|---|---|---|
| DK710 | sarA-gp130_Y759F_Fwd | Y759F in codon-opt *sarA:gp130* | 5'-gcaccacggtggagaactgcactgtgcta-3' |
| DK711 | sarA-gp130_Y759F_Rvr | Y759F in codon-opt *sarA:gp130* | 5'-tagcacagtgcagttctccaccgtggtgc-3' |
| DK793 | sarA_A159Y_H161T_N162V_Fwd | ASHN → YSTV in codon-opt *sarA* | ctgagcaatgtagccgcttggggga**AC**a**GT**gct**ATA**gggtgtggtacaactagtagagcc |
| DK794 | sarA_A159Y_H161T_N162V_Rvr | ASHN → YSTV in codon-opt *sarA* | ggctctactagttgtaccacaccc**TAT**agc**AC**t**GT**tcccccaagcggctacattgctcag |
| DK1127 | FLAG_SarA_pY-1_R_fwd | I168R in codon-opt *sarA* | gccgatactgagccctgtagccgcttgggggа |
| DK1128 | FLAG_SarA_pY-1_R_rev | I168R in codon-opt *sarA* | tcccccaagcggctacagggctcagtatcggc |
| DK1129 | FLAG_SarA_pY-2_H_fwd | A169H in codon-opt *sarA* | gtgccgatactgatgaatgtagccgcttggggga |
| DK1130 | FLAG_SarA_pY-2_H_rev | A169H in codon-opt *sarA* | tcccccaagcggctacattcatcagtatcggcac |
| DK1131 | FLAG_SarA_pY-1-2_RH_fwd | Y**IA**Q → Y**RH**Q in codon-opt *sarA* | gcgctgtgccgatactgatgcctgtagccgcttgggggattatg |
| DK1132 | FLAG_SarA_pY-1-2_RH_rev | Y**IA**Q → Y**RH**Q in codon-opt *sarA* | cataatcccccaagcggctacaggcatcagtatcggcacagcgc |
| DK1133 | FLAG_SarA_pY-1_L_fwd | I168L in codon-opt *sarA* | ccgatactgagcaaggtagccgcttgggg |
| DK1134 | FLAG-SarA_pY-1_L_rev | I168L in codon-opt *sarA* | ccccaagcggctaccttgctcagtatcgg |
| DK1135 | FLAG_SarA:gp130_pY-1_I_fwd | R168I in codon-opt *sarA:gp130* | cggagggcacctggtgtatatagccagagtgcaccac |
| DK1136 | FLAG_SarA:gp130_pY-1_I_rev | R168I in codon-opt *sarA:gp130* | gtggtgcactctggctatatacaccaggtgccctccg |
| DK1020 | sarA-gp130_R168I_H169A_F | Y**RH**Q → Y**IA**Q in codon-opt *sarA :gp130* | ggagggcacctgggctatatagccagagtgcaccacggtggag |
| DK1021 | sarA-gp130_R168I_H169A_R | Y**RH**Q → Y**IA**Q in codon-opt *sarA :gp130* | ctccaccgtggtgcactctggctatatagcccaggtgccctcc |
| DK1141 | psk129_sarA_t503g_c504g_fwd | I168R in native *sarA* | atgcctgtattgagccctataaccggacggtgggttatg |
| DK1142 | psk129_sarA_t503g_c504g_rev | I168R in native *sarA* | cataacccaccgtccggttatagggctcaatacaggcat |
| DK1143 | pwsk129_sarA_g505c_c506a_fwd | A169H in native *sarA* | gcagaatgcctgtattgatggatataaccggacggtgg |
| DK1144 | pwsk129_sarA_g505c_c506a_rev | A169H in native *sarA* | ccaccgtccggttatatccatcaatacaggcattctgc |
| DK1145 | pwsk129_sarA:gp130_c502a_g503t_fwd | Y**RH**Q → Y**IA**Q in *sarA :gp130* | ggaacctggtgaatataaccgctgtgaaccacggtgc |
| DK1146 | pwsk129_sarA:gp130_c502a_g503t_rev | Y**RH**Q → Y**IA**Q in *sarA :gp130* | gcaccgtggttcacagcggttatattcaccaggttcc |

| | | | |
|---|---|---|---|
| DK1170 | FLAG-gp130_R238I_fwd | R238I in codon-opt *gp130dimer* | 5'-ggcacctggtgtatatagccggagtgcaccacg-3' |
| DK1171 | FLAG-gp130_R238I_rev | R238I in codon-opt *gp130dimer* | 5'-cgtggtgcactccggctatatacaccaggtgcc-3' |
| Trx651 | SarAΔN20_fwd | Clamp, PciI, SarAΔN20 | cgcgggacatgtca GATGTTAATTTAGAGGAC |
| IOP197 | SarAΔN20_I168R_rev | Clamp, NotI, stop codon, SarA with I68R mutation | cgcgggGCGGCCGCTTATTCATCCGGGAAAACCTCTGCAGAATGCCTGTATTGAGCGCGATAACCGGACGGTGG |

### Table S2: Primers

| Strain | Genotype | Plasmid | Resistance | Derived From | Notes |
|---|---|---|---|---|---|
| DCK22 | 14028s | p67GFP | Amp | CS093 | |
| DCK444 | 14028s Δ*sarA* | P67GFP | Amp | DCK440 | |
| DCK487b | 14028s Δ*sarA* | pWSK129-*sarA*, p67GFP | Amp, Kan | DCK444 | |
| DCK1156 | 14028s Δ*sarA* | pWSK129-*sarA:gp130*, p67GFP | Amp, Kan | DCK869 | |
| DCK1222 | 14028s Δ*sarA* | pWSK129-*sarA*$^{I168R}$, p67GFP | Amp, Kan | DCK1224 | |
| DCK1223 | 14028s Δ*sarA* | pWSK129-*sarA:gp130*$^{R168I}$, p67GFP | Amp, Kan | DCK1225 | |

#### Table S3: Bacterial Strains

Braun, D.A., Fribourg, M., and Sealfon, S.C. (2013). Cytokine response is determined by duration of receptor and signal transducers and activators of transcription 3 (STAT3) activation. J Biol Chem *288*, 2986-2993.

Jaslow, S.L., Gibbs, K.D., Fricke, W.F., Wang, L., Pittman, K.J., Mammel, M.K., Thaden, J.T., Fowler, V.G., Jr., Hammer, G.E., Elfenbein, J.R., *et al.* (2018). Salmonella Activation of STAT3 Signaling by SarA Effector Promotes Intracellular Replication and Production of IL-10. Cell Rep *23*, 3525-3536.
